# Development of itaconate polymer microparticles for intracellular regulation of pro-inflammatory macrophage activation

**DOI:** 10.1101/2025.01.30.635692

**Authors:** Kaitlyn E. Woodworth, Zachary S.C.S. Froom, Natasha D. Osborne, Christian N. Rempe, Brenden Wheeler, Kyle Medd, Neal I. Callaghan, Huikang Qian, Abhinav P. Acharya, Carlie Charron, Locke Davenport Huyer

## Abstract

Itaconate (IA) is an endogenous metabolite and a potent regulator of the innate immune system. Its use in immunomodulatory therapies has faced limitations due to inherent challenges in achieving controlled delivery and requirements for high extracellular concentrations to achieve internalization of the highly polar small molecule to achieve its intracellular therapeutic activity. Microparticle (MP)-based delivery strategies are a promising approach for intracellular delivery of small molecule metabolites through macrophage phagocytosis and subsequent intracellular polymer degradation-based delivery. Toward the goal of intracellular delivery of IA, degradable polyester polymer-(poly(itaconate-co-dodecanediol)) based IA polymer microparticles (IA-MPs) were generated using an emulsion method, forming micron-scale (∼ 1.5 µm) degradable microspheres. IA-MPs were characterized with respect to their material properties and IA release kinetics to inform particle fabrication. Treatment of murine bone marrow-derived macrophages with an optimized particle concentration of 0.1 mg/million cells enabled phagocytosis-mediated internalization and low levels of cytotoxicity. Flow cytometry demonstrated IA-MP-specific regulation of IA-sensitive inflammatory targets. Metabolic analyses demonstrated that IA-MP internalization inhibited oxidative metabolism and induced glycolytic reliance, consistent with the established mechanism of IA-associated inhibition of succinate dehydrogenase. This development of IA-based polymer microparticles provides a basis for additional innovative metabolite-based microparticle drug delivery systems for the treatment of inflammatory disease.

## 1 Introduction

Chronic inflammation is a key driving factor in the pathology of many non-communicable diseases (NCDs),^[1]^ where altered regulation of initiation, maintenance, and resolution of inflammation defines disease pathology.^[2]^ ^[3]^ Macrophages embody a central role in inflammatory control, capable of initiating inflammation via the secretion of pro-inflammatory cytokines and chemokines in response to pathogens, and conversely, regulating the pathogenesis of inflammation by transitioning their phenotype.^[4]^ Given the central role of macrophages in NCDs, current immunomodulation strategies often involve drug or cytokine-mediated targeting of macrophages to directly modulate immune-related signaling processes. However, the effectiveness of local and systemic immunomodulatory therapies is challenged by adverse effects, including toxicity and non-specific immunosuppression. These issues are partly a consequence of the need for precise cellular targeting, the complexity of fine-tuning macrophage phenotypes owing to the broad pharmaceutical mechanisms, and ineffective therapeutic delivery to specific tissues.^[5]^ There is a need for novel specific strategies to control macrophage-directed inflammation.[6].

In response to inflammatory stimuli, such as lipopolysaccharide (LPS) and type I/II interferons (IFN), macrophages undergo metabolic reprogramming. Upon stimulation, succinate dehydrogenase (SDH) becomes active, driving the production of reactive oxygen species (ROS) and stabilizing hypoxia-inducible factor (HIF)-1α.^[7]^ This stabilization promotes the production of pro-inflammatory interleukin (IL)-1β and influences the expression of other inflammation-associated genes.^[8]^ Concurrently, the tricarboxylic acid (TCA) cycle flux decreases, forcing the cell to rely more heavily on glycolysis to meet its energy demands.^[9]^ During this inflammatory state, itaconate (IA), a small carboxylic acid metabolite and an intrinsic regulator of inflammation, begins to accumulate.^[10]^ IA plays a critical role in counteracting inflammation by directly inhibiting SDH, which halts succinate oxidation, leading to a cessation of ROS production, destabilization of HIF-1α, and suppression of IL-1β production.^[11]^ Subsequent accumulation of succinate marks a significant metabolic shift within the cell.^[9]^ IA further dampens the inflammatory metabolic profile by inhibiting key glycolytic enzymes through cysteine modifications,^[12]^ reducing the reliance on glycolysis and enhancing TCA cycle activity through glutamate oxidation.^[13]^ IA also contributes to the regulation of oxidative stress as it stabilizes erythroid 2-related factor 2 (Nrf2), which mitigates oxidative damage and regulates the electrophilic stress response. Additionally, IA modulates inflammatory signaling by influencing NF-κB activity and reduces the expression of IL-6 through an Nrf2-independent mechanism involving the IκBζ−ATF3 axis.^[14]^

The established mechanisms of IA as an intrinsic immunomodulator highlights its potential as a biomimetic therapy in chronic inflammatory conditions. Unfortunately, the polarity of IA makes it challenging to passively diffuse through cell membranes, necessitating specific transport mechanisms or an innovative delivery system. The immunomodulatory potential of IA has been investigated using membrane-permeable IA derivatives, such as dimethyl itaconate (DMI) and 4-octyl itaconate (4OI), which have demonstrated their ability to inhibit the expression of pro-inflammatory mediators like IL-6, IL-12, TNF-α and IL-1β in LPS-stimulated macrophages.^[15]^ However, exogenous IA derivatives do not recapitulate the full electrophilic stress response and transcriptional profile of endogenous IA.^[16]^ Furthermore, the administration of IA derivatives does not enhance intracellular IA levels,^[17]^ whereas non-modified IA accumulates in both resting and activated macrophages.^[16]^ This suggests that unmodified IA has a distinct advantage compared to its electrophilic derivatives.

The bioavailability of IA remains a challenge, especially when administered orally, as it is quickly removed from circulation;^[18]^ IA exerts the bulk of its therapeutic activity at millimolar concentrations, a level that is difficult to maintain either systemically or locally for a relevant timespan. To overcome these challenges, we previously developed polymers which degrade over time *in situ* to release IA at therapeutic concentrations.^[19]^ This approach provides extended anti-inflammatory effects with a high loading capacity (50% of the material mass), and controlled release kinetics (i.e., days-years) imparted through material chemistry approaches.^[20]^

Phagocytosis is an intricate process enabling macrophages and other innate immune cells to engulf and degrade pathogens. The phagocytic ability of macrophages offers an opportunity for selective metabolite-based immunomodulatory therapy given an appropriate vehicle,^[6]^ providing intracellular delivery tools targeted to inflammatory phagocytes. Polymeric microparticle drug delivery systems have been widely used toward this goal, leveraging tunable material properties including size and release kinetics^[21]^ to deliver combinations of antigen, adjuvant, chemokines, and other immunomodulating molecules through phagocytosis and subsequent controlled release. The intracellular degradation rate of these microspheres can be regulated by altering the molecular weight and monomer composition of the copolymers that make up the microspheres.^[22–24]^ Notably, numerous microparticle intracellular delivery technologies have been published with respect to poly(lactide-*co*-glycolide) (PLGA)-based delivery vehicles to achieve a variety of delivery applications.^[25–34]^ For example, intracellular delivery of dexamethasone-loaded MPs can successfully downregulate inflammatory gene expression in human macrophages over several weeks, while minimizing systemic effects of prolonged glucocorticoid administration.^[5,35]^ In dendritic cells, phagocytotic delivery of α-ketoglutarate polymer particle technologies have demonstrated sustained metabolite release, vaccine adjuvant delivery, and T-cell regulation in inflammatory models. ^[36–44]^

In this work, we describe the optimization of IA-based polymer technologies for the development of polymer microparticles (IA-MP) to act as a potential therapeutic strategy to target pro-inflammatory macrophage activity with IA (**Figure 1A**). We introduce a robust and scalable method for synthesizing IA-based polyester microparticles (IA-MPs) designed for controlled intracellular degradation and release of IA. IA-MPs effectively modulate macrophage function, mimicking the immunoregulatory effects and metabolic rewiring effects of endogenous IA. Given the challenge of delivering IA systemically due to its limited potency and uptake, this platform offers a promising alternative for localized immune modulation.

**Figure 1:**
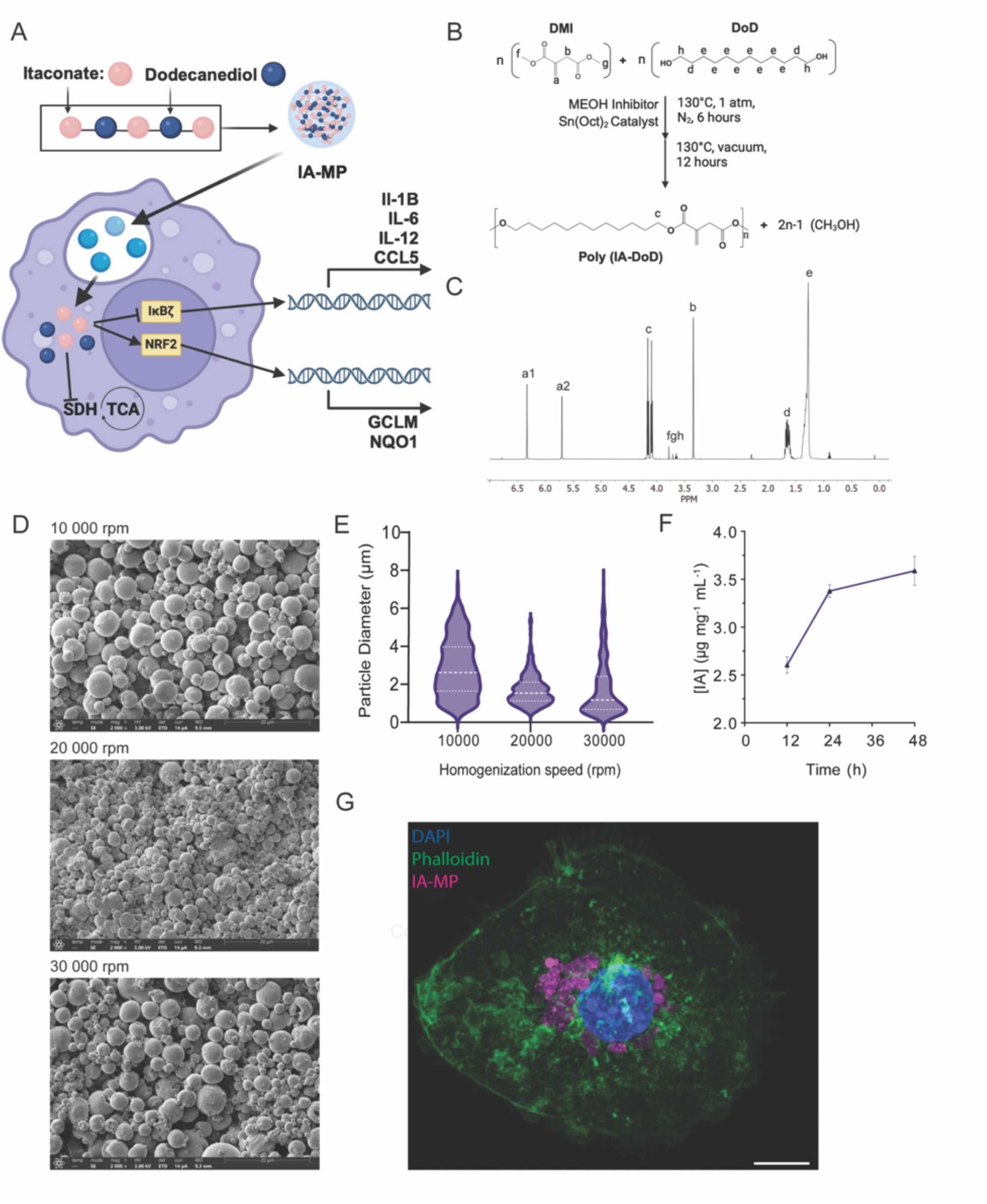
Development of degradable itaconate-containing polymer microparticles. A) Fabrication of polymer microparticles provides an intracellular delivery method of itaconate to modulate macrophage metabolic function and regulate inflammatory pathways. B) Bulk polycondensation of DMI and DoD in the presence of a radical inhibitor and a tin-based catalyst generates an IA polymer. C) Polymer purity and structural characterization using 1H NMR. D) Scanning electron microscope image of IA-MPs generated using increasing homogenization speeds (scale bar = 20 µm), and E) corresponding violin plot of IA-MP size distributions. F) Quantified soluble IA released from hydrolytically degraded IA-DoD polymer in Milli-Q water over 48 h (n = 2). G) Confocal image of internalized IA-MP stained with DAPI (blue; cell nuclei); Rhodamine 6G (magenta; labelled IA-MPs), and Phalloidin 647 (green, F-actin) (scale bar = 10 µm).

## 2 Materials and methods

### 2.1 Materials

DMI, deuterated chloroform (CDCl3), tin (II) 2-ethylhexanoate, Rhodamine 6G, polyvinyl alcohol (PVA, MW 89,000–98,000, 99+% hydrolyzed), LPS, 4OI, bovine serum albumin (BSA), dimethyl sulfoxide (DMSO), formic acid (FA), acetonitrile (ACN) and puromycin were purchased from Sigma Aldrich (St. Louis, MO). Anti-puromycin (clone 12D10, AF647) was purchased from Millipore Sigma (Burlington, MA). Dichloromethane (DCM) was purchased from VWR Chemicals (Mississauga, ON). Recombinant human M-CSF was purchased from VWR Chemicals (Radnor, PA). Lymphoprep sucrose gradient was purchased from StemCell Technologies (Vancouver, BC). 1,12-Dodecanediol (DoD), Invitogen ebioscience Fluoromount G, Hoechst 33342, SYTOX Green Nucleic Acid Stain, DAPI, 4% Paraformaldehyde (PFA), and PCR primers (*Canx, Rer-1, Il-1β, Il-6, Nos2, Hif1α, Nqo1, Gclm, Gsr Taldo, Hmox1, Glut-1, HK-1, SDHA, PKM-1, Ccl5, Il-12p40*) were purchased from Thermo Scientific (Waltham, MA). 4-methoxyphenol (MEHQ) was purchased from RCI America (Edison, NJ). RPMI 1640 Medium with L-glutamine, Fetal Bovine Serum (FBS), Dulbecco’s phosphate buffered saline (DPBS), penicillin/streptomycin solution, ethylenediaminetetraacetic acid (EDTA), and Trypsin-EDTA (0.05%) were purchased from Corning (Corning, NY). ACK lysis buffer was purchased from Quality Biological (Gaithersburg, MD). Macrophage colony-stimulating factor (M-CSF), Zombie NIR Fixable Dye, TruStain FcX, monocyte blocker, anti-CD86 (clone GL-1, BV421), FOXP3 fixation/permeabilization solutions, anti-TNFα (clone MP6-XT22, BV605), anti-IL12 p40 (clone C15.6, PE-Cy7), and anti-F4/80 (clone BM8, BV605) were purchased from BioLegend (San Diego, CA). Itaconic acid (≥99.0%) and 2-deoxyglucose (2-DG) were purchased from TCI America (Portland, OR). Chloroform, tetrahydrofuran (THF), and TRIzol were purchased from Fisher Scientific (Waltham, MA). SuperScript IV VILO Master Mix was purchased from Invitrogen (Carlsbad, CA). SsoAdvanced Universal SYBR® Green Supermix was purchased from Bio-Rad Laboratories (Hercules, CA). Anti-MHC-II (clone M5/114.15.2, BV786) and anti-IL6 (clone MP5-20F3, APC) were purchased from BD Biosciences (Franklin Lakes, NJ). Oligomycin A, harringtonine, and Phalloidin-iFluor 647 were purchased from Abcam (Cambridge, UK). RNeasy Plus Mini Kit and DNase set were purchased from Qiagen (Toronto, ON). EasiVial polystyrene low standard was purchased from Agilent Technologies (Santa Clara, CA). All materials were used as received unless otherwise described.

### 2.2 Polymer synthesis and characterization

Poly(itaconate-*co-*dodecanediol) (poly(IA-DoD)), was synthesized using a one-pot, two-stage melt condensation technique using a 1:1.1 molar ratio of DMI and DoD (10 g basis of DMI). The catalyst, tin(II) 2-ethylhexanoate, was added at 0.01 mol of catalyst per mol of ester, and MEHQ, an inhibitor of radical polymerization, was added at 0.5 weight percent of all reagents. The reaction proceeded for 6 hours at 130°C under a nitrogen atmosphere, followed by 18 hours under vacuum (<5 kPa), with constant mechanical stirring (200 rpm). The polymer was subjected to two rounds of precipitation in -80°C methanol to remove catalyst, short chain oligomers, and unreacted monomers, then dried at room temperature for 48 hours ^[19]^.

Polymer structure was confirmed using proton nuclear magnetic spectroscopy (^1^H-NMR). Samples were dissolved in deuterated CDCl3 at a concentration of approximately 10 mg mL^-1^, and recorded on a Bruker Avance NEO operating at a frequency of 500 MHz. Chemical shifts (δ) were reported in parts per million (ppm) relative to the residual proton signal of CDCl3 at 7.26 ppm. Peak integration was performed to calculate percent esterification based on the following formulas:

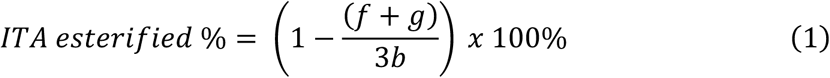

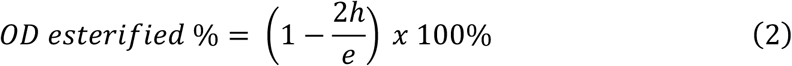

Polymer molecular weight was analyzed via gel permeation chromatography (GPC) using an Agilent 1260 Infinity Multi-Detector system fitted with a PLgel 7.5 x 50 mm, 10 µm, guard column (Agilent Technologies, Santa Clara, CA) and two PLgel 5 µm mixed-D columns (Agilent Technologies, Santa Clara, CA). Samples were dissolved in THF (2 mg mL^-1^), and passed through a 0.22 μm polytetrafluoroethylene (PTFE) filters prior to addition to sample vials. All replicates were run at a eluent flow rate of 1 mL min^-1^ at 50°C. Calibration was performed using EasiVial polystyrene low standard (51950 – 162 MW range) and a 28.9 kDa nominal Mp standard. Results were analyzed on Agilent GPC-SEC Software (v.2.8) (Agilent Technologies, Santa Clara, CA) using RI conventional GPC analysis to determine the number-average molecular weight (Mn), weight-average molecular weight (Mw), and dispersity (Đ = Mw/Mn). Melting point was determined by packing the solid polymer into a glass capillary and measurement using a Mel-Temp apparatus equipped with a mercury thermometer.

### 2.3 Quantification of degradation release of itaconate

Soluble release of IA from poly(IA-DoD) was quantified following incubation of poly(IA-DoD) particulate (1 g mL^-1^) in MilliQ water (15 MΩ cm^-1^) under agitation (300-400 rpm) for up to 48 h (12, 24, and 48 h samples). Supernatant was collected, flash frozen in liquid nitrogen, and stored at -80°C before further analysis. Degradation supernatant was subsequently warmed to room temperature, vigorously vortexed to resolubilize any precipitate, and passed through a 0.2 µm PTFE filter to remove any insoluble particulate. Quantification of IA release was conducted via UV absorbance at 264 nm via high-performance liquid chromatography (HPLC) with metabolite confirmation by tandem mass spectrometry (MS). Calibration curves for accurate quantification of IA were modeled using prepared standards at concentrations ranging from 0.01 mmol L^-1^ to 10 mmol L^-1^. All samples and standards were run on an Agilent LC/MSD equipped with an Agilent C-18 ZORBAX reverse phase column and appropriate guard column at 1.5 mL min^-1^ using Milli-Q water and ACN (both containing 0.1% (v/v) FA) as solvents. The gradient was run over 10 min from 2-30% (v/v) ACN. Prior to each sample injection set, an IA standard at 5 mmol L^-1^ was submitted to control for between-run variability in the performance of the HPLC. Using Agilent OpenLab Data Analysis software (v.2.7) (Agilent Technologies, Santa Clara, CA), IA was quantitated by the area under the curve of the 264 nm absorbance elution peak.

### 2.4 Microparticle synthesis and characterization

The poly(IA-DoD) polymer microparticles (poly(IA-DoD) MPs) were synthesized using a water-in-oil-in-water (w/o/w) emulsion method ^[22]^. Poly(IA-DoD) polymer material (50 mg) was dissolved in 4 mL of DCM, followed by homogenization (Fisherbrand^TM^ Homogenizer Model 1000) with 60 mL of 2% (w/v) PVA solution for 3 minutes at 20 000 rpm to create the primary emulsion. Particles were also fabricated at 10 000 and 30 000 rpm to assess size changes with speed of homogenization. An additional 40 mL of 1% PVA was added to the primary emulsion and stirred overnight to allow for organic solvent evaporation. Poly(IA-DoD) MPs were concentrated using centrifugation (10 min, 10 000 x g), washed (3X) with deionized distilled water, flash frozen in liquid nitrogen and lyophilized (24 h, - 60°C, 57 kPa) for storage (-20°C) for future experiments. For experimentation where visualization of particles with fluorescence was required, 4 mg of Rhodamine 6G was added to the initial DCM solution prior to homogenization, and the same process was followed. Characterization of IA-MP structure was assessed using scanning electron microscopy (SEM). IA-MPs were resuspended in Milli-Q water (15 MΩ cm^-1^), pipetted onto glass slides, and left to dry overnight in a fume hood. Dried material from the slides was attached to SEM stubs with carbon tape for gold sputter-coating. SEM images were obtained using a Hitachi S-4700 FE Scanning Electron Microscope. Particle diameters were manually measured using ImageJ (US National Institutes of Health, Bethesda, MD) ^[45]^.

### 2.5 Bone marrow-derived monocyte isolation and culture

All animal experiments were performed in accordance with the Canadian Council on Animal Care (CCAC) and approved by the University Committee on Laboratory Animals of Dalhousie University (protocol # 22-026). Bone marrow was isolated from the hindlimbs of C57BL/6 mice (male and female, Charles River Laboratories, 6-12 weeks old). Animals were euthanized with isoflurane and carbon dioxide gas before hind limbs were removed and collected in complete RPMI media (RPMI 1640 with 2 mmol L^-1^ L-glutamine, 10% FBS, 1% penicillin/streptomycin). Bones were debrided of soft tissue, rinsed in 70% ethanol, washed with DPBS, then sectioned and flushed with complete RPMI media to collect bone marrow. Bone marrow was homogenized using a pestle, filtered through a 40 µm cell strainer, concentrated (centrifuged at 300 x g, 5 min), and treated with ACK lysis buffer (1 min at RT, 1 mL mouse^-1^) to remove red blood cells prior to differentiation. Cells were plated at 10 million cells per 8.8 cm-diameter plate (60.8 cm^2^) in complete RPMI media (10 mL) and differentiated to bone marrow derived macrophages (BMDMs) by addition of 20 ng mL^-1^ of murine M-CSF for 7 days, with 50% media changes on day 3 and 5.

### 2.6 Particle toxicity analysis

Particle associated temporal changes in cell number and endpoint toxicity was tested using BMDMs and characterized through temporal live cell imaging using a BioTek Cytation 1 and BioSpa 8 automated incubator (Agilent Technologies, Santa Clara, CA) fitted with an automated laser autofocus cube, DAPI (EX 377/50 nm, EM 447/60 nm, dichroic mirror 409 nm, LED 365 nm) and GFP (EX 469/35 nm, EM 525/39 nm, dichroic mirror 497 nm, LED 465 nm) filter cubes. Cells were treated for 24 h at 50 000 cells well^-1^ in a 96 well plate. BMDM were stained with Hoechst 33342 (1 μg mL^-1^, 10 min) then subsequently treated with a serial dilution of poly(IA-DoD) particles (1, 0.5, 0.25, 0.1, 0.05, 0.025, 0.01, 0.005, 0.001 mg mL^-1^) prepared in complete RPMI media with M-CSF (20 ng mL^-1^) and LPS (100 ng mL^-1^), or a particle free media control. Wells were imaged every 6 hours to quantify changes in cell count. At 24 h, supernatant was removed, cells were washed (3X) with DPBS and stained with SYTOX green nucleic acid stain (1 μmol L^-1^, 10 min). Imaging was performed following toxicity staining to capture cell number and associated death. Imaging analysis was performed using Agilent BioTek Gen5 Software (v.2.07). Cell counts calculated in BioTek Gen5 using primary masking of the nucleus (Hoechst 33342) in the 447 nm channel. Secondary masking in the 525 nm channel (SYTOX) was constrained to the primary mask and thresholds were set based on untreated cells respectively to generate dead and live cell counts. Cytotoxicity was then calculated as percent cell death using the dead count compared to the live count and overall cell count using BioTek Gen5 (Agilent Technologies, Santa Clara, CA).

### 2.7 Confocal imaging

Particle internalization was performed using confocal imaging. Cells were treated on coverslips for 12 h at 500 000 cells per well in a 12 well plate with 0.1 mg mL^-1^ of IA-MP prepared in complete RPMI media with M-CSF (20 ng mL^-1^) and LPS (100 ng mL^-1^). At 12h, cell culture media was removed, BMDMs were washed (3X) with DPBS, and fixed for 10 minutes with 4% PFA in DPBS. The cells were washed with DPBS (3X) and permeabilized with 0.1% tween (10 min) before subsequent washing staining with DAPI (1 μg mL^-1^, 10 minutes), and Phalloidin-iFluor 647 (1X, 60 min). Coverslips were slide mounted with 20 μL of Fluoromount G and stored in the dark at -20°C until imaging. Confocal Z stacks were obtained using a ZEISS LSM880 (Oberkochen, Germany) inverted scope at 63X oil, NA 1.4 (Plan-Apochromat lens) magnification. Maximum intensity projections of Z stacks were generated using Zen 3.6 microscopy software (ZEISS, Oberkochen, Germany).

### 2.8 Gene expression

IA-MP regulation of macrophage inflammation was assessed using BMDMs (1 million cells • 9.6 cm^-2^) treated for 24 h with 100 ng mL^-1^ of LPS, alone or simultaneously with IA-MPs (0.1 mg million cells^-1^), IA (5 mmol L^-1^), or 4OI (0.125 mmol L^-1^). To yield a population of viable cells that contained IA-MP, the IA-MP (Rhodamine 6G labelled) treated group were trypsinized, scraped, and pelleted at 300 x g for 5 min, and transferred to a 96-well round-bottom polystyrene plate for staining. Cells were stained with Zombie NIR Fixable Dye (1:1000 dilution in DPBS) for 30 minutes at 4°C in the dark and washed twice with 1X DPBS solution containing 1% (w/v) bovine serum albumin (BSA) and 1 mmol L^-1^ EDTA (FACS Buffer). Surface antibody staining (45min, 4°C) was performed with anti-F4/80 (1:187.5) in FACS buffer containing TruStain FcX (1:50) and Monocyte Blocker (1:50) (Surface Staining Buffer), cells were then washed twice (FACS buffer), and resuspended in FACS Buffer for fluorescence activated cell sorting. Viable IA-MP containing macrophages (ZombieNIR-, F4/80+, Rhodamine 6G+ ) were collected on a BD FACS Aria Fusion^TM^ (BD Biosciences, Franklin Lakes, New Jersey, USA), and lysed in TRIzol (750 μL). In the other treatment groups (IA, 4OI, LPS alone), adherent cells were cell scraped and lysed in 750 μL of TRIzol per well. All lysate was stored at -80°C until further RNA extraction.

RNA extraction was performed through addition of chloroform (150 μL) to each thawed tube of Trizol lysate. After centrifugation (12 000 x g, 15 min, 4°C), the clear aqueous RNA containing phase was transferred for RNA purification using the Qiagen RNeasy Plus Mini Kit (Qiagen, Hilden, Germany) according to the manufacturer’s instructions. Purified RNA was subsequently quantified using the μDrop plate (Thermo Scientific, Waltham, MA) in a Varioskan LUX Multimode Microplate Reader (Thermo Scientific, Waltham, MA). cDNA was synthesized using SuperScript IV VILO Master Mix as per manufacturer’s directions using Applied Biosystems 2720 Thermal Cycler (Thermo Scientific, Waltham, MA). Real-time quantitative polymerase chain reaction (RT-qPCR) was used to measure gene expression using CFX384 Real Time System C1000 Touch Thermocycler (Bio-Rad Laboratories, Hercules, CA). Using the *Mus musculus* transcriptome (Mouse GRCm39) annotated by the Ensembl genome browser (version 113, European Bioinformatics Institute, Cambridgeshire, UK), primers were designed to be exon-spanning, and they were confirmed to be transcript-specific and to not bind gDNA by using NCBI Primer-BLAST (version 2.16.0, National Library of Medicine, Bethesda, MD).^[46,47]^ IDT’s online OligoAnalyzer tool (Coralville, IA) was used to confirm no secondary structures were predicted (including primer dimers) and the optimal annealing temperature (Ta) of each primer was determined by Ta Gradient. Primers for genes of interest (0.20 µM working concentration, per reaction; summarized in **Table S1**) were combined with SsoAdvanced Universal SYBR® Green Supermix (5 µL per reaction), 3 µL cDNA (5 ng per reaction) and adjusted to a total volume of 10 µL per reaction with nuclease-free water. Each sample was run in triplicate and the raw Ct values were assessed using Bio-Rad CFX Maestro software (version 1.1 Bio-Rad Laboratories, Hercules, CA) and exported for further analysis. All genes of interest were analyzed using the 2^-ΔΔCt^ method, using difference in Ct of the gene of interest from the geometric mean of three stable housekeeping genes (*Stx5a*, Syntaxin 5A; *Calnx*, Calnexin; and *Rer1*, Retention In Endoplasmic Reticulum Sorting Receptor 1), normalized to the LPS treated alone control sample group, to yield Relative Gene Expression.

### 2.9 Flow cytometry staining and analysis

BMDMs (1 million cells / 9.6 cm^2^) were treated for 12 h with 100 ng mL^-1^ of LPS, alone or simultaneously with Rhodamine 6G labelled IA-MPs (0.1 mg million cells^-1^), soluble IA (5 mmol L^-1^), or 4OI (0.125 mmol L^-1^). Cells were trypsinized, scraped, and pelleted at 300 x g for 5 min, and transferred to a 96-well round-bottom polystyrene plate for staining. Cells were stained with Zombie NIR Fixable Dye (1:1000 dilution in DPBS, 30min, 4°C), washed twice with FACS Buffer, then surface stained (45min, 4°C) with anti-CD86 (1:200) and anti-MHC-II (1:500) in Surface Staining Buffer. Cells were then washed twice with FACS buffer, and then fixed and permeabilized using the Biolegend FOXP3 fixation/permeabilization kit in accordance with manufacturer protocols. Intracellular staining of anti-TNFα (1:200), anti-IL6 (1:200) and anti-IL12 p40 (1:500 was performed in FOXP3 permeabilization buffer with TruStain FcX (1:50) for 30 min at room temperature. Cells were then washed with FACS and resuspended in 1X DPBS for collection on the cytometer. Flow cytometry data was collected using a BD FACSCelesta^TM^ (BD Biosciences, Franklin Lakes, New Jersey, USA), and analysis was conducted in FlowJo v.10.9.1, (BD Biosciences, Franklin Lakes, New Jersey) (**Figure S1, Table S2**), with fluorescence-minus-one controls (FMOs) prepared for each marker.

### 2.10 Single-cell functional metabolic profiling

Live samples were profiled in single-cell format for metabolic function by flow cytometry using methods adapted from the validated SCENITH workflow.^[48]^ Briefly, protein synthesis (measured by exogenous puromycin incorporation) correlates to whole-cell metabolic rate. Single-cell puromycin content is assessed across differential application of specific metabolic inhibitors to determine relative metabolic pathway dependencies of cells positively identified with concurrent standard flow cytometric expressional profiling. Samples were divided into four groups and plated in a polystyrene tissue-culture plate at 1 x 10^6^ cells per well. After 24 h of stimulation (100 ng mL^-1^ LPS), for each sample, each group received a media change (1 mL RPMI 1640 with 10% FBS and 1% P/S) containing a metabolic inhibitor. For each sample, aliquots were incubated with oligomycin A (ATP synthase inhibitor; 1 µmol L^-1^) 2-DG (glycolytic inhibitor; 100 mmol L^-1^), harringtonine (negative control for protein synthesis; 2 μg mL^-1^), or DMSO (vehicle control; 1:1000 v/v) at 37°C for 30 min. Puromycin (protein synthesis marker; 10 µg mL^-1^) was then introduced and samples were incubated for an additional 30 minutes. Samples were washed with DPBS and analyzed by flow cytometry as previously described. Cells were surface stained with anti-F4/80 (1:375) and anti-MHC-II (1:500), then stained intracellularly with anti-puromycin (1:1000). Data was collected on a BD Celesta cytometer and analyzed with FlowJo v.10.9.1 **(Figure S2, Table S3)**.

### 2.11 Macrophage respirometry

Blood was obtained from three female donors (IRB under Hematopoietic Biorepository and Cellular Therapy, Case Western Reserve University), and Lymphoprep sucrose gradient was utilized to isolate peripheral blood mononuclear cells (PBMCs). These PBMCs were then differentiated into macrophages by culturing the cells the presence of 50 ng mL^-1^ recombinant human M-CSF for 7 days. Macrophages were were seeded at 50 000 cells well^-1^ in Seahorse™ XFe96 cell culture plates (Agilent Technologies, Santa Clara, CA) for a mitochondrial stress test according to manufacturer directions. Seahorse™ XF base medium was supplemented with (in mmol L^-1^) glucose (10), L-glutamine (2), and sodium pyruvate (1). Oxygen consumption rates (OCR) were measured 4 times per successive treatment (in µmol L^-1^) of neat medium, oligomycin (3), FCCP (1), and rotenone (1). OCRs were averaged across within-treatment timepoints, except for in the FCCP treatment where the single peak OCR was used.

### 2.12 Statistical methods

All data were first assessed for normality (Anderson–Darling, D’Agostino, Shapiro–Wilk, and Kolmogorov–Smirnov tests) and homogeneity of variances (Brown-Forsythe and Bartlett’s tests), and parametric or non-parametric statistics were performed as appropriate, using GraphPad Prism (version10.0, Dotmatics, La Jolla, CA, USA). Two-way ANOVA followed by pairwise comparisons with Tukey’s multiple comparisons test method were used to determine the statistical significance and assess the interactive effects of factors in Figure 2B and 4C. One-way ANOVA followed by pairwise comparisons with Tukey’s multiple comparisons test method (normal distribution), or a non-parametric Kruskal-Wallis test paired with a Dunn test were used to determine the statistical significance and assess the interactive effects of factors in Figure 2A, 3, and 4E-G.

**Figure 2:**
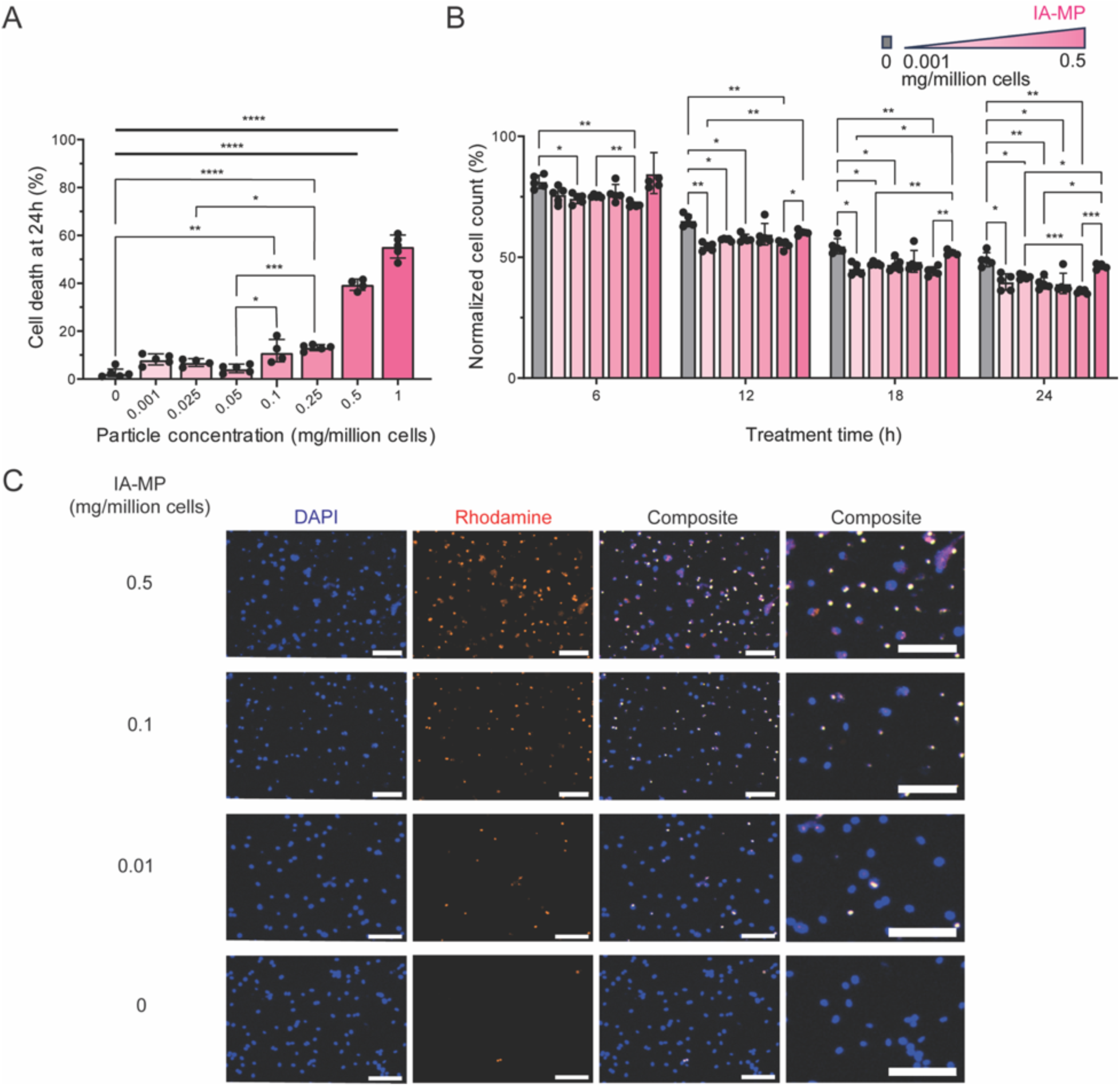
IA-MPs are internalized and minimally toxic in LPS-treated bone marrow-derived monocytes. Cell death (%, A) and total cell count (%, B) after 24 h IA-MP treatment (IA-MP concentrations: 0, 0.001, 0.025, 0.05, 0.1, 0.25, 0.5 mg/million cells). C) Representative BMDM uptake (DAPI (blue; cell nuclei); Rhodamine 6G (red; labelled IA-MPs)) after 24 h of IA-MP treatment of 0.5, 0.1, 0.01, and 0 mg/million cells (scale bar = 100 µm). Data are expressed as mean ± SD, n = 6. Statistical significance is indicated as * p < 0.05, ** p < 0.01, *** p < 0.001, **** p < 0.0001.

## 3 Results

### 3.1 Generation of IA polymer microparticles for intracellular delivery

Phagocytosable microparticles were designed for intracellular degradation and metabolite release, based on previously developed polyester IA-based materials ^[19]^ and first optimized for molecular weight (MW) and melting temperature (**Figure 1B**). Synthesis of poly(IA-DoD) through bulk condensation maintained the pendent unsaturated carbonyl group, which is critical for IA’s immunoregulatory capacity. Poly(IA-DoD) was a solid material at body temperature (37°C; melting temperature: 50°C), suggesting its applicability for particle development. Structure assessment of poly(IA-DoD) polymer material using ^1^H-NMR and subsequent peak identification confirmed the formation of a linear polymer material with maintenance of the pendant unsaturated carbonyl on IA that drives much of the immunoregulatory capacity (**Figure 1C**). Quantification of polymer end groups through peak integration indicated an esterification of IA and DoD of 96.57% and 100% respectively (Formulas 1-2), suggesting IA end capped the generated polymer chains. The number average molecular weight (Mn) of poly(IA-DoD) was 10,860 ± 1,190 g/mol with a Đ of 2.26 ± 0.15.

Using a water-in-oil-in-water emulsion method, we generated spherical polymer microparticles from poly(IA-DoD) material (**Figure 1D**). To target the biologically active range of macrophage phagocytosis, particles were designed to be within 0.1-10 μm in diameter.^[49]^ IA-MP particle size was inversely related to homogenization speed, with median MP diameters of 2.6 µm (10 000 rpm), 1.5 µm (20 000 rpm) and 1.2 µm (30 000 rpm) **(Figure 1E)**, suggesting potential benefit for future manipulation of particle properties. IA release in polymer supernatant treated with deionized distilled water was quantified and shown to increase over 48 h (**Figure 1F**).

### 3.2 IA-MPs are internalized and non-cytotoxic

IA-MPs were readily internalized by macrophages, visualized by confocal microscopy and counter membrane staining with phalloidin (**Figure 1G**). Live cell imaging of macrophages exposed to IA-based polymeric microparticles at varying concentrations (0.001, 0.01, 0.025, 0.05, 0.1, 0.25, and 0.5 mg • 10^6^ cells^-1^) was conducted to evaluate particle phagocytosis, cell viability, and apoptotic behavior at 24 h of treatment (**Figure 2**). Quantitative cell death staining revealed significant increases in cell death in treatments with 0.5 and 1 mg of IA-MP • 10^6^ cells^-1^ compared to lower particle concentrations and the LPS media control (**Figure 2A)**, indicating a threshold beyond which IA-MP loading induces cytotoxic effects. Cell counts achieved through nuclear labelling and serial quantification normalized to initial values showed significant decreases in particle-treated groups relative to the LPS media control (**Figure 2B**), however percent cell death remained below 20% in all wells up to an IA-MP concentration of 0.25 mg/million cells. Representative endpoint images at concentrations of 0, 0.01, 0.1, and 0.5 mg • 10^6^ cells^-1^ confirmed this trend of increased cell loss with higher particle concentrations (**Figure 2C**). After 24 hours of treatment with 0.1 mg IA-MP • 10^6^ cells^-1^, 99.96 ± 0.05% of BMDMs exhibited successful particle internalization (**Figure S3**); demonstrating that efficient uptake can be achieved without substantial induction of cell death.

### 3.3 Macrophage immunomodulation by IA-based microparticle degradation

RT-qPCR analysis was conducted to assess the expression of NF-κB and Nrf2 target genes in LPS-stimulated BMDMs. NF-κB-dependent genes, including *IL-6*, *Ccl5*, and *IL-12p40*, exhibited a significant reduction in expression in the IA-MP-treated group compared to the LPS control. This reduction was consistent with that observed in cells treated with soluble IA at mM concentration (5 mmol L^-1^) described elsewhere,^[50,51]^ extending to other genes including *IL-1β*, *Nos2*, and *Hif1α* (**Figure 3A**). In contrast, Nrf2-regulated genes did not exhibit significant changes in expression levels in the IA-MP-treated macrophages relative to the LPS-stimulated controls, but a trend in upregulated expression was observed that matched soluble IA treatment (**Figure 3A**).

**Figure 3:**
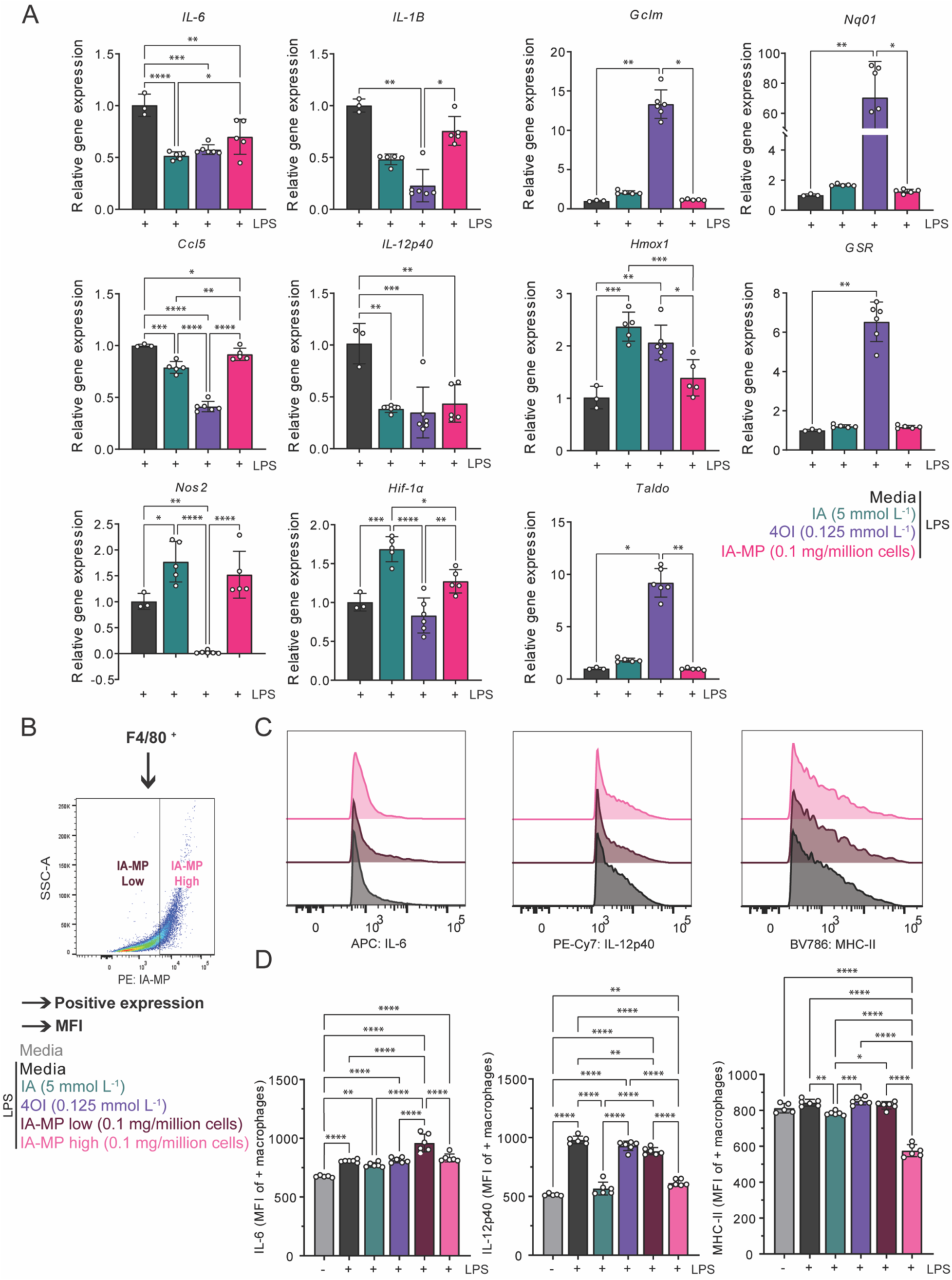
IA-MPs modulate inflammatory macrophage signaling in an uptake-specific manner. Comparative BMDM expression of pro-inflammatory macrophage markers. A) Relative transcript expression (2^-ΔΔCT^) of *IL-6*, *IL-1β*, *Ccl5*, *IL-12p40*, *Nos2*, *Hif1α*, *Gclm*, *Nqo1*, *Hmox1*, *Gsr*, and *Taldo*. B-D) Expression of pro-inflammatory markers by IA-MP-treated BMDMs is altered by IA-MP uptake. B) IA-MP-treated BMDMs were gated according to PE (Rhodamine 6G) and SSC to generate low- and high-uptake groups for analysis. C) Representative histograms of IL-6, IL-12p40, and MHC-II expression in IA-MP low- and high-uptake subpopulations. D) Quantitative expression of IL-6, IL-12p40, and MHC-II in macrophage-based treatment with IA (5 mmol L^-1^), 4OI (0.125 mmol L-1), or IA-MPs (low and high relative uptake). Data are expressed as mean ± SD, n = 3 (media) or 6 (experimental treatments). Statistical significance is indicated by * p < 0.05 ** p < 0.01, *** p < 0.001, **** p < 0.0001.

Flow cytometric analysis was undertaken to quantify macrophage IA-MP uptake and characterize the resultant cell phenotype. To fully appreciate IA-MP loading specific functionality, we isolated cells positive for expression of IL-6, IL-12p40, and MHC-II (12h and 24h (**Figure S4**)), then subsequently gated macrophages based on absolute internalization of IA-MP determined by PE intensity (Rhodamine 6G signal from particles) vs. side scatter signal (SSC-A) (**Figure S3**). As nearly 100% of IA-MP macrophages were positive for particle uptake at this concentration, we next analyzed cells based on *high* and *low* relative particle uptake based on Rhodamine 6G expression and shifts in granularity (SSC-A) (**Figure 3B**). Median fluorescence intensity (MFI) in the positive population of IL-12p40, and MHC-II (**Figure 3C-D, Figure S4B**) were significantly reduced in the IA-MP high-uptake group when compared to IA-MP low-uptake loaded cells and particle-free LPS stimulated controls. IL-6 MFI was also reduced in the IA-MP high-uptake group compared to IA-MP low-uptake samples.

### 3.4 Macrophage metabolic dependencies respond to IA-MP administration

The SCENITH approach to characterization of metabolic utilization was used to explore changes to metabolic organization associated with IA-MP uptake (**Figure 4**). This method measures protein synthesis and corresponding exogenous puromycin incorporation as a proxy for whole-cell metabolic rate in the presence of the metabolic inhibitors 2-DG (a glycolytic inhibitor) or oligomycin A (ATP synthase inhibitor) (**Figure 4A**). IA-MP-treated BMDMs displayed a significantly higher puromycin MFI under oligomycin treatment compared to LPS-stimulated and unstimulated controls, indicating reduced dependence on oxidative phosphorylation (**Figure 4B-C**). Correspondingly, 2DG decreased puromycin MFI significantly in the same treatment, further indicating primarily glycolytic dependence.

**Figure 4:**
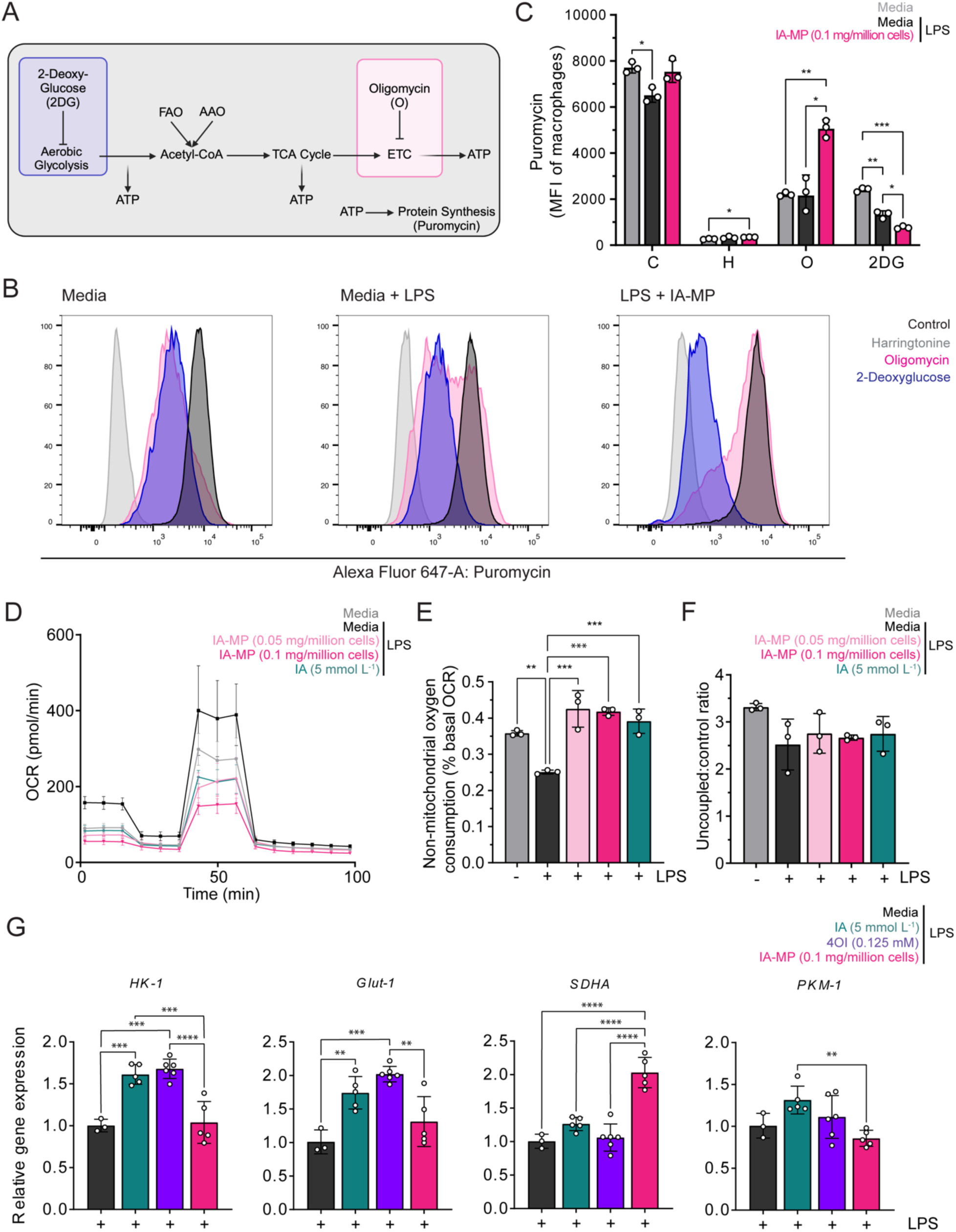
IA-MPs alter macrophage metabolic function. A) Graphic overview of flow cytometric SCENITH method used to assess reliance on oxidative or glycolytic metabolism through quantification of protein synthesis (corresponding to level of puromycin incorporation). B) Modal histogram representation of SCENITH-quantified puromycin levels differentiated by control (black), harringtonine (grey), oligomycin (pink), and 2DG (purple) treatment conditions. C) SCENITH-quantified puromycin mean fluorescence intensity in control, harringtonine, oligomycin, and 2DG conditions of macrophages treated with IA-MPs with respect to media +/- LPS. D-F) Respirometric outputs of human macrophage cultures in control, low- or high-dose IA-MPs, or bulk IA treatments, including D) oxygen consumption rate, E) non-mitochondrial oxygen consumption, and F) uncoupled:control ratio. G) Relative transcript expression (2^-ΔΔCT^) of Glut-1, HK-1, SDHA, and PKM-1. Data are expressed as mean ± SD, n = 3 (media) or 6 (experimental treatments). Statistical significance is indicated as *p < 0.05 ** p < 0.01, *** p < 0.001, **** p < 0.0001.

Respirometric analysis of peripheral blood derived human macrophages (**Figure 4D**) revealed significantly higher normalized non-mitochondrial oxygen consumption in the IA-MP treated group (**Figure 4E**), but no significant difference in peak oxidative capacity (**Figure 4F**) in IA-MP-treated groups relative to the control condition, mirroring trends observed with soluble IA (5 mmol L^-1^).

Despite these metabolic shifts, gene expression analysis of key glycolytic markers (*Glut-1*, Glucose transporter 1; *HK-1*, Hexokinase 1, *PKM-1,* pyruvate kinase M1) showed a slight upregulation of *Glut-1* and no significant differences in *HK-1* or *PKM-1* expression between BMDMs treated with IA-MP and LPS-stimulated controls (**Figure 4G**). Interestingly, IA-MP treatment led to a significant upregulation of *SDHA* compared to both the LPS control and cells treated with soluble IA or 4OI at concentrations described elsewhere.^[14,50–53]^

## 4 Discussion

IA has garnered considerable attention as an effective regulator of innate immunity, making it a compelling candidate for the development of anti-inflammatory therapeutics. However, effective delivery of IA has proven challenging to realize with traditional systemic approaches based on its low absolute potency and barriers to uptake.^[54,55]^ This limitation extends to its derivatives, such as 4OI, which are membrane-permeable but do not fully recapitulate the endogenous effects of IA.^[16,56]^ Here, we have demonstrated the development of degradable IA-based polymer microparticles (IA-MPs) that can achieve sustained and controlled delivery of IA directly to macrophages based on their phagocytic activity, overcoming some of the challenges associated with systemic administration. IA-MPs offer tunability as a drug delivery system, as particle size, degradation rates, and IA release can be adjusted based on the synthesis and processing conditions,^[57]^ making them a promising platform for local inflammatory control.

Phagocytic BMDMs readily internalized IA-MPs while cell mortality remained low even at higher MP concentrations, suggesting a technology comparable to other MP delivery approaches described elsewhere.^[58,59]^ In experiments probing BMDM function, phagocytosed IA-MPs regulated cytokine production and expression of antigen presentation-associated proteins. Gene expression analysis revealed significant regulation of the NRF2 and NF-κΒ pathway genes in IA-MP-treated macrophages comparable to 5 mmol L^-1^ IA, a previously demonstrated efficacious concentration of exogenous IA administration.^[50,51]^ The specific regulation of these pathways, which are crucial for cellular stress responses and inflammation, supports the role of IA-MPs in achieving IA-specific regulation at a transcriptional level. This highlights the precise immunomodulatory effects facilitated by the microparticle delivery system, ensuring control over macrophage activity in inflammatory conditions without the requirement for millimolar extracellular IA concentrations. IA-MP-mediated regulation of BMDM phenotype was dependent on the degree of particle uptake; flow cytometric analysis revealed that macrophages with higher internalization of IA particles exhibited reduced levels of inflammatory IL-6, IL-12p40, and MHC-II expression. This suggests IA release by IA-MP degradation shifts macrophages toward an anti-inflammatory state,^[60]^ consistent with increased localized and intracellular IA release that is increased higher levels of IA-MP phagocytosis.^[50]^ MHC-II, which plays a key role in antigen presentation and immune activation, displayed reduced expression in macrophages with higher particle uptake, suggesting a shift toward a more tolerogenic phenotype. This finding has important implications for the use of IA-MPs in conditions where immune regulation is desirable, such as chronic inflammation or autoimmunity, where controlled downregulation of antigen presentation may help mitigate excessive immune responses and support tolerogenic responses.^[61]^

Mechanistically, IA-mediated inhibition of SDH and its downstream effects on metabolic reprogramming are central to its anti-inflammatory properties. By disrupting the TCA cycle and reducing succinate-driven ROS production, IA dampens pro-inflammatory signaling and promotes a metabolic shift toward glycolysis.^[62]^ Metabolic analyses recapitulated this metabolic shift in IA-MP-treated macrophages, demonstrating reduced oxidative phosphorylation and increased reliance on glycolysis, consistent with marked inhibition of SDH and resultingly lower TCA flux. Interestingly, although robustly enhanced by LPS stimulation, the transcript abundance of key glycolysis-associated genes *Glut-1* and *Hk-1* exhibited either no or minor changes based on the presence or concentration of IA-MPs. However, because functional utilization indicated increased glycolytic dependence in IA treated groups, taken together these results suggest that the metabolic regulation induced by IA-MP treatment is largely either on an expression or post-translational level.

While the development of IA-MPs represents a promising approach for phagocyte specific intracellular immunomodulation, several limitations in this study highlight the need for further investigation. *In vitro* models do not fully capture the complexity of *in vivo* immune responses and long-term therapeutic effects. Specifically, the extended LPS stimulation used in BMDM cultures can lead to an exhausted macrophage phenotype,^[63]^ limiting the ability to assess the full spectrum of macrophage behavior over extended periods. Additionally, the immune microenvironment *in vitro* is simplified, lacking the dynamic interactions between immune cells, extracellular matrix components, and other factors that influence the response to therapeutic agents *in vivo*. Consequently, although the in *vitro* data offer valuable insights into IA-MP uptake and initial metabolic reprogramming, they lack the capacity to fully capture the intricacies of chronic inflammation or other disease states. An *in vivo* environment presents additional complexities including the presence of diverse enzymes, location-specific microenvironments, and variations in localized water dynamics, all of which can influence MP degradation and therapeutic performance ^[64–66]^. *In vitro*, the repeated exposure of macrophages to IA-MPs may not accurately replicate the sustained release and degradation dynamics observed *in vivo*, where particle retention and gradual degradation over time are essential for maintaining therapeutic efficacy. The optimization of polymer properties, including the degradation profile and release kinetics, will be necessary to ensure that the IA-MPs provide controlled and sustained delivery of the therapeutic payload.

The demonstration of IA-MP-enabled uptake and selective metabolic regulation in activated phagocytes suggests an immunoregulatory platform that bypasses the traditional challenges of IA-based therapeutic candidates. The benefit of a degrading delivery vehicle that concomitantly serves as the active therapeutic, enables optimization of delivery latency, sustained release, release rate, or the co-incorporation of complementary therapeutics. Future directions include the demonstration of functional immunomodulatory benefit *in situ* in models of chronic inflammation, such as the foreign body response or local lesions of autoimmune etiology.

## 5 Data availability

Data will be made available on request.

## 6 Acknowledgements

This research is supported by the Dalhousie Medical Research Foundation Faculty of Dentistry Early Career Research Award, Natural Sciences and Engineering Research Council of Canada (NSERC) Discovery Grant (RGPIN-2022-03666), NSERC Alliance International Catalyst Grant (ALLRP 597098-24), New Frontiers in Research Fund – Exploration Fund (NFRFE-2022-00313), and Canadian Institutes for Health Research Project Grant (PJT – 191692), NIAID, NIH, USA award R01AI155907 and NIAMS, NIH, USA award R01AR078343. We acknowledge infrastructure support from the Canadian Foundation for Innovation (JELF 42294, JELF 39824), Research Nova Scotia Research Opportunities Fund (2022-2400, 2020-1208), and NSERC Research Tools and Instrumentation Grant (RTI-2024-00128, RTI-2017-00320). KW, ZF, CR, and BW were supported by a Canada Graduate Scholarship – Masters. KW, ZF, and BW were Nova Scotia Graduate Scholarship, ZF and BW were supported by a Killam Predoctoral Scholarship. BW was supported by a Canada Graduate Scholarship – Doctoral. The authors would like to acknowledge the assistance and expert advice from Dr. Penghao Xiao at Dalhousie University’s Department of Chemistry Mass Spectrometry Core Facility.

## 7 Author contributions

KW conceived the idea, designed the experiments, analyzed the results, and wrote the manuscript. ZF assisted with in vitro experiments and performed live cell imaging and analysis. NO performed RT-qPCR and analyzed the results. CR designed and performed the SCENITH experiment. BW completed the degradation experiment, and performed molecular weight characterization and analysis. KM ran HPLC and analyzed the results. NC assisted in writing the manuscript. HQ and APA performed Seahorse respirometry. CC advised on mass spectrometry assessment. LDH supervised the entire project. All authors read and edited the manuscript and approved its final submission.

## 9 Appendix. Supplementary Materials

Supplementary material includes Table S1, Table S2, Table S3, Figure S1, Figure S2, Figure S3, and Figure S4.

