## Supplementary Tables and Figures for "Development of itaconate polymer microparticles for intracellular regulation of pro-inflammatory macrophage activation"

.

### 1 Supplementary Tables

**Table S1:** Murine RT-qPCR primers.

| Gene name | Forward primer sequence (5' to 3') | Reverse primer sequence (5' to 3') |
|---|---|---|
| *Canx* | ATGGAAGGGAAGTGGTTACTGT | GCTTTGTAGGTGACCTTTGGAG |
| *Rer1* | GGCTGGACAAGTCTACCCC | GCGTAGGTCACAATGTACCAAC |
| *Stx5a* | GGGTCCATCATGTCAATAG | GGGTCCATCATGTCAATAG |
| *IL-1β* | AGCAGCACATCAACAAGAGC | GGAAAAGAAGGTGCTCATGTCC |
| *IL-6* | TTCTTGGGACTGATGCTGGTG | CAGAATTGCCATTGCACAACTC |
| *Nos2* | TTTGTGCGAAGTGTCAGTGG | TTGTGCTGGGAGTCATGGAG |
| *Hif1α* | CAAAGTTCACCTGAGAGACTTC | TGGATTCTTTGCCTCTGTGTC |
| *Nqo1* | TTCTGTGGCTTCCAGGTCTT | TCCAGACGTTTCTTCCATCC) |
| *Gclm* | ACAATGACCCGAAAGAACTGC | GAGTACCTCAGCAGCCACAG |
| *Gsr* | GCGTGAATGTTGGATGTGTACC | GTTGCATAGCCGTGGATAATTTC |
| *Taldo* | CGGGTGATTTCAATGCCATCG | GCCTCCTCTACCAGCTCTTG |
| *Hmox1* | AGGTACACATCCAAGCCGAGA | CATCACCAGCTTAAAGCCTTCT |
| *HK-1* | CCATGCGGCTCTCTGATG | GGACAAAGGTTGGCAGCATC |
| *Glut-1* | GCTGTGCTTATGGGCTTCTC | CACATACATGGGCACAAAGC |
| *SDHA* | GCAGCACAGGGAGGTATC | CTAGCTCAACCACAGAGG |
| *PKM-1* | CAGCCATGTTCCACCGTC | TGAGCACTCCTGCCAGAC |
| *Ccl5* | AGGACTCTGAGACAGCAC | CATATGGTGAGGCAGGTG |

| | | |
|---|---|---|
| *Il-12p40* | TGTGGAATGGCGTCTCTGTC | GCGGGTCTGGTTTGATGATG |

**Table S2:** Flow cytometry panel design for cytokine and expression marker analysis.

| Fluorophore | Antigen | Clone |
|---|---|---|
| BV421 | CD86 | GL-1 |
| Zombie AQUA | Viability | |
| BV605 | TNF-α | MP6-XT22 |
| BV786 | MHC-II | M5/114.15.2 |
| PE | IA-MP | |
| PE-Cy7 | Il-12/Il-23 p 40 | C15.6 |
| APC | Il-6 | MP5-20F3 |
| APC750 | iNOS | CXNFT |

**Table S3:** Flow cytometric panel design for SCENITH analysis.

| Fluorophore | Antigen | Clone |
|---|---|---|
| BV421 | CD86 | GL-1 |
| BV605 | F4/80 | BM8 |
| BV786 | MHC-II | M5/114.15.2 |
| PE | IA-MP | |
| Alexa Fluor 647 | Puromycin | 12D10 |
| APC-Cy7 | Viability | |

#### 2 Supplementary Figures

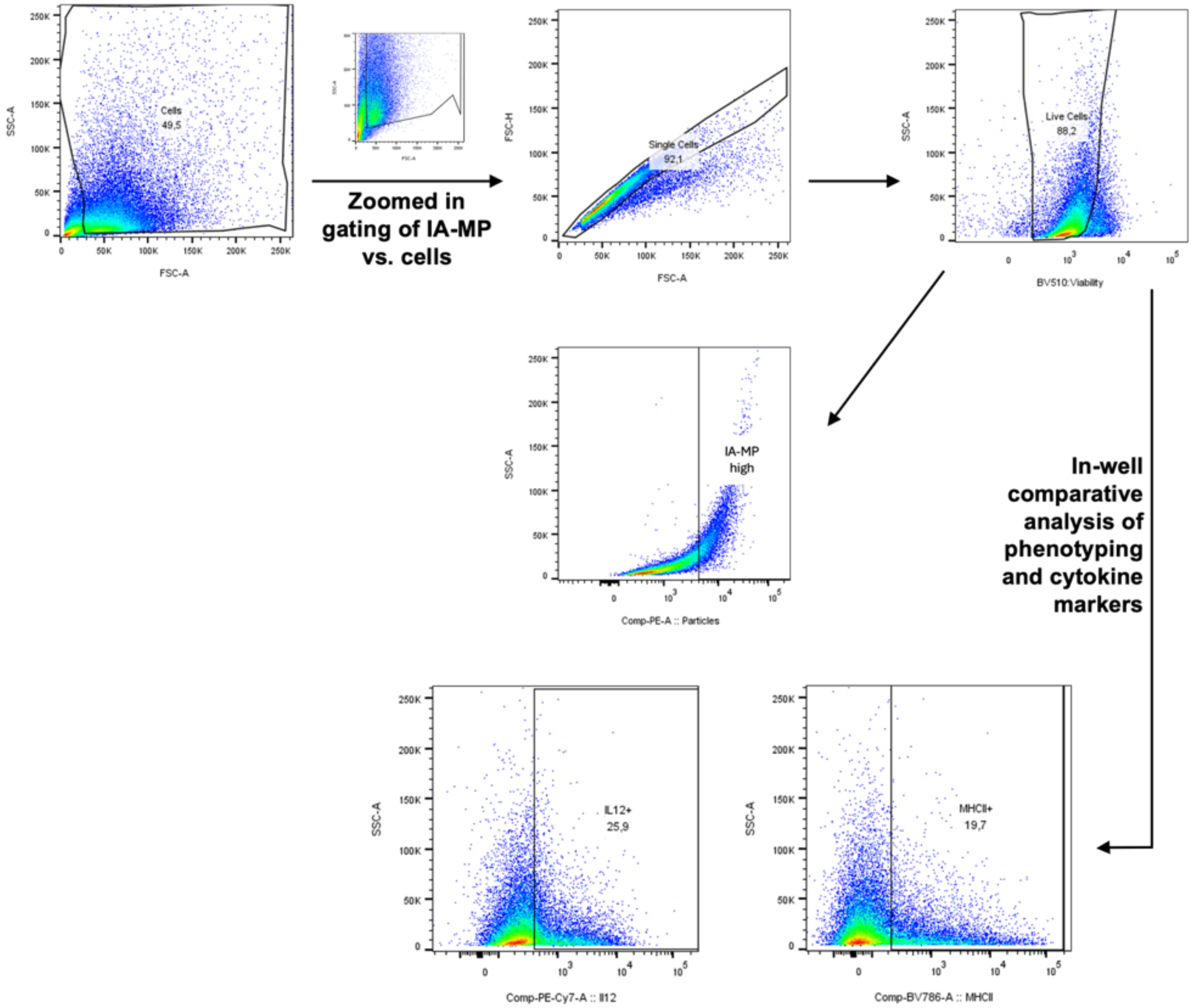


**Figure S1:** Flow cytometry gating scheme for cytokine and expression marker analysis.

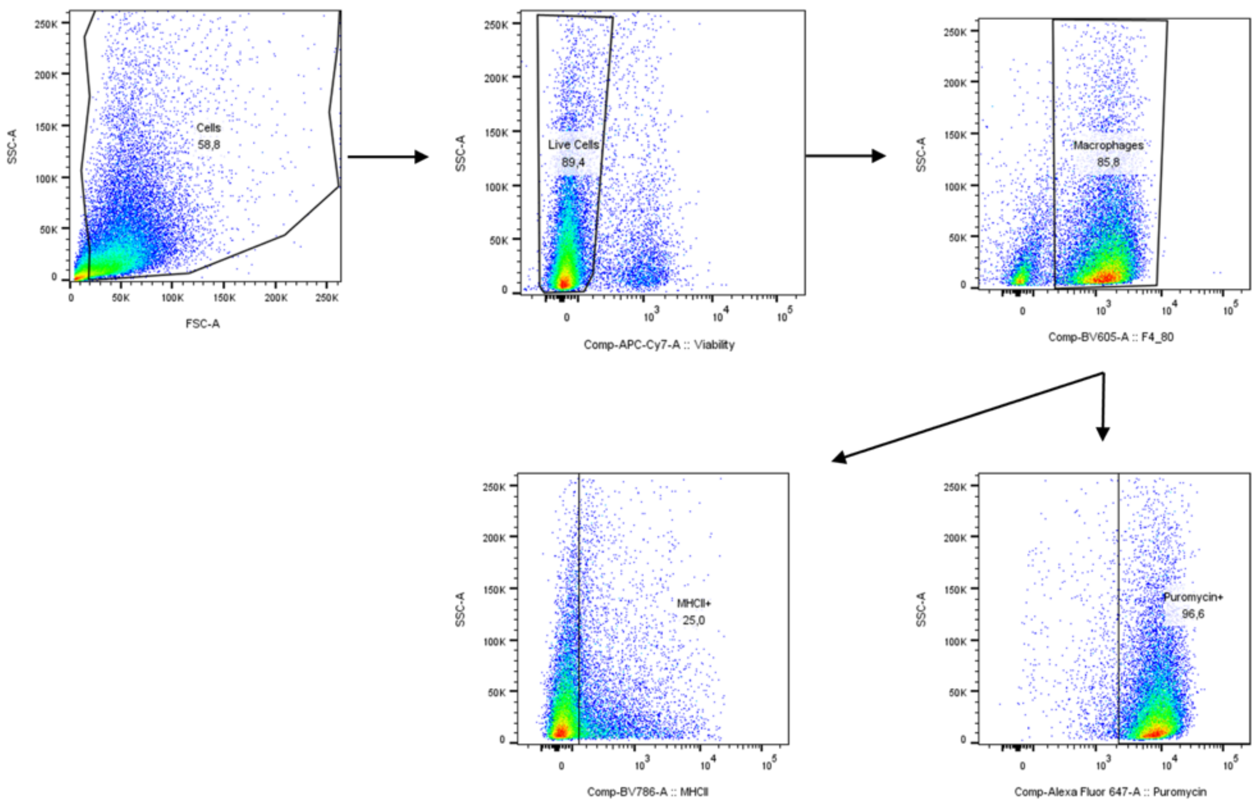


**Figure S2:** Flow cytometric gating scheme for SCENITH analysis.

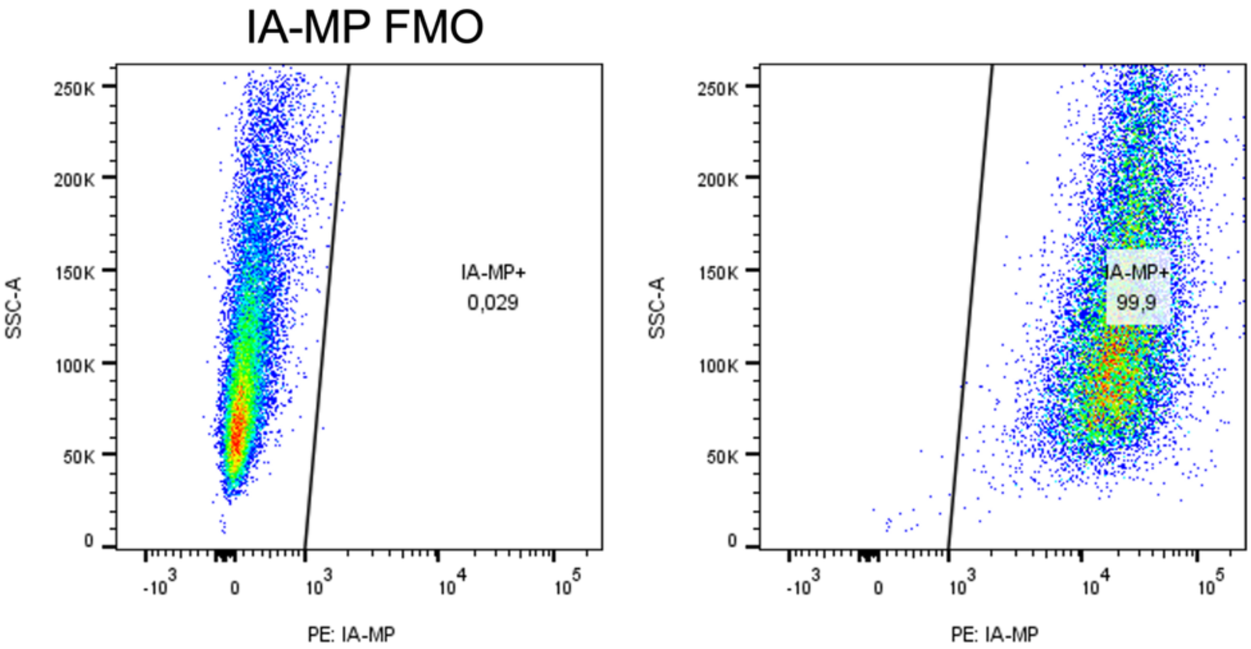


**Figure S3:** Flow cytometric IA-MP gating for uptake quantification using IA-MP FMO. IA-MP FMO on left, IA-MP treatment condition on right showing 99.9% uptake in the sample.

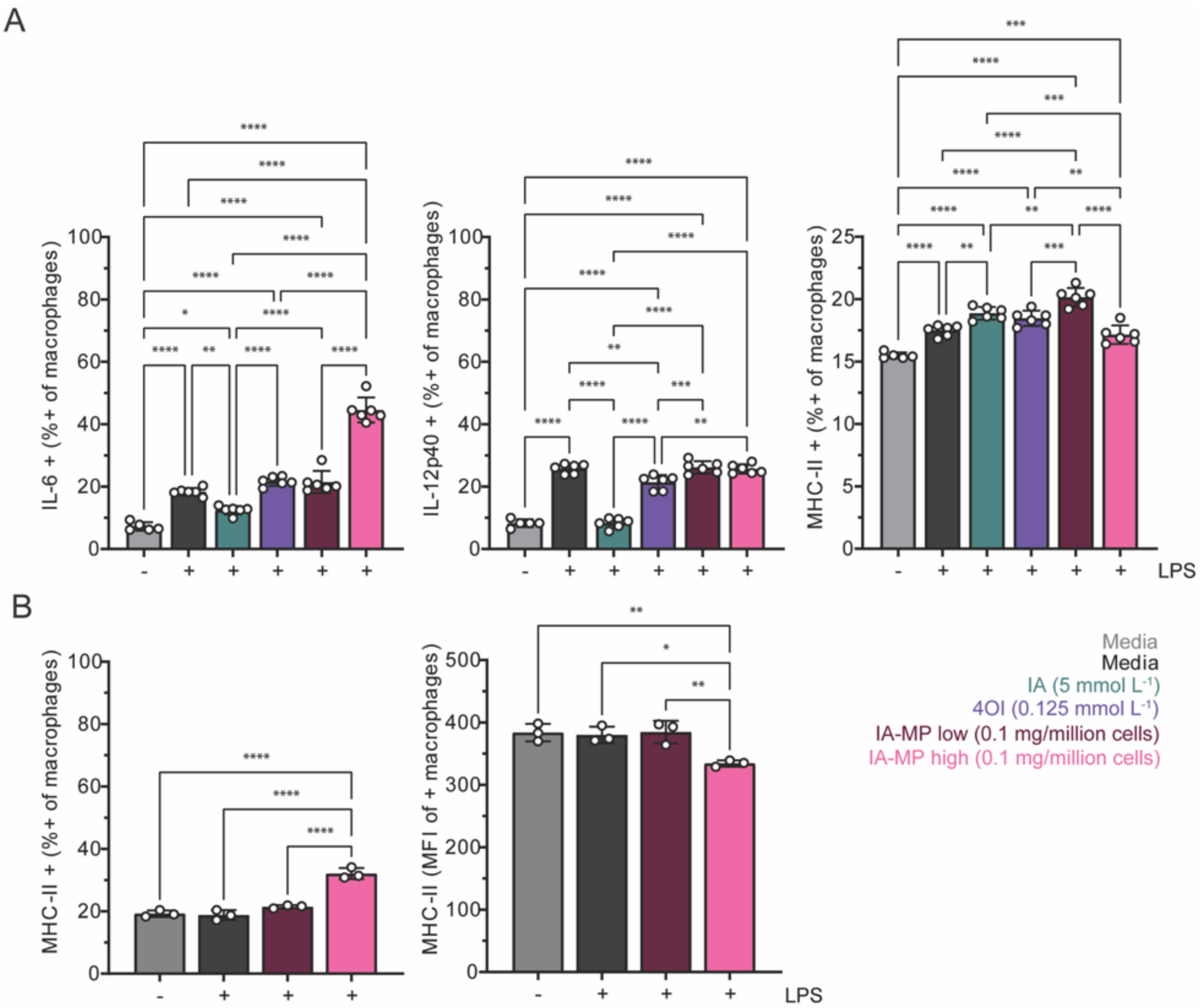


**Figure S4:** A) Percent positive populations of Il-6, Il-12p40, and MHC-II with IA-MP high- and low- uptake subpopulations. B) Percent positive and subsequent MFI of the positive population for MHC-II at 24h. Data are expressed as mean ± SD, n = 3 (B) or 6 (A). Statistical significance is indicated as *p < 0.05 ** p < 0.01, *** p < 0.001, **** p < 0.0001.
